# ANGPTL6 variant induces cerebral vascular dysfunction and predisposes to intracranial aneurysm in mice

**DOI:** 10.1101/2025.01.10.632391

**Authors:** Céline Baron-Menguy, Milène Freneau, Mary-Adel Mrad, Corentin Lebot, Marc Rio, Vincent L’allinec, Thibaut Quillard, Hubert Desal, Richard Redon, Romain Bourcier, Anne-Clémence Vion, Gervaise Loirand

## Abstract

**Background:** Intracranial aneurysm (IA) is a common cerebrovascular abnormality characterized by localized dilation and wall thinning in intracranial arteries, that frequently leads to fatal sub-arachnoid hemorrhage. Pathophysiological mechanisms responsible for AI remains largely unknown, but increasing evidence suggest that genetic susceptibility plays a predominant role. We recently identified a rare nonsense variant in *ANGPTL6* gene that prevented angiopoietin-like 6 (ANGPTL6) secretion and predisposed to IA. The aim of this study is now to understand why the ANGPTL6 variant predisposes to IA.

**Methods:** *Angptl6*-knock in mice were generated by homologous recombination. Cerebral arteries of the circle of Willis have been analyzed under basal and hemodynamic overload conditions. Functional properties of cerebral arteries have been analyzed by pressure arteriography. Effect of recombinant ANGPTL6 have been assessed on vascular smooth muscle cells and ANGPTL6 partners have been analyzed by surface plasmon resonance.

**Results:** *Angptl6*-knock in mice display endothelial dysfunction expressed by reduced NO production in response to flow in cerebral arteries. They present exaggerated dilatation of cerebral arteries under hypertensive stress, and aggravated wall damage and aneurysmal remodeling of the intima and media at arterial bifurcations of the circle of Willis under hemodynamic overload. Matrix bound ANGPTL6 decreased VSMC migration and increased adhesion and FAK signaling activation, without affecting VSMC phenotype. Surface plasmon resonance analyses identified αvβ5 integrin as a new ANPTL6 receptor.

**Conclusions:** *Angptl6*-knock in mice mimicked the main features of IA in humans. ANGPTL6 appears to be an extracellular matrix component regulating vascular cell function, which may be involved in mechanosensing and generating adapted vascular cell responses to hemodynamic stress. Non-secreted ANGPTL6 variants would result in the loss of this function, thus favoring IA under conditions of hemodynamic overload.

## Introduction

Intracranial aneurysm (IA) is a frequent and generally asymptomatic cerebrovascular abnormality affecting 3-5% of the general population.^1^ IA is a localized dilation and wall thinning of intracranial arteries preferentially occurring at the bifurcations of the circle of Willis, where hemodynamic stress is high. The unpredictable rupture of IA leads to subarachnoid hemorrhage with high morbidity.^2,3^ Besides the surgical or endovascular interventions commonly used to treat ruptured IAs, or to prevent IA rupture in at risk patients,^4^ there are no therapeutic options, particularly pharmacological ones, to manage this life-threatening conditions. Understanding the pathophysiological mechanisms of initiation, progression and rupture of IAs that may allow the identification of therapeutic targets of interest remains challenging. The fact that the disease is limited to a restricted vascular area with difficult access prevents the development and use of relevant cellular models. Furthermore, the difficulty of generating reliable experimental models that reproduce the pathogenesis of the disease^5^ considerably hinders the discovery of pathophysiological mechanisms that their use could allow.

Increasing evidence suggest that genetic factors contribute to disease susceptibility.^6^ Identifying these factors might be very useful to better understand the physiopathology of IAs, and potentially to predict their formation and rupture.^7,8^ However, identification of specific genes or causal molecular pathways by genome-wide association studies remains largely inconclusive and these common alleles together explain only a small fraction of the risk attributable to genetics for IA in the general population.^9–12^ More recently, whole-exome sequencing which can detect rare coding variants that have large effect and induce a high risk of developing IA have proven to be effective in providing valuable insights into the pathophysiology of IA and pave the way for new diagnostic or therapeutic strategies.^13–19^ Thanks to such an approach, we recently we identified rare nonsense variants in *ANGPTL6* encoding angiopoietin-like 6 (ANGPTL6) in families affected with IA.^14^ This discovery of *ANGPTL6* variants in families with multiple IA carries was then replicated in two independent studies, thus supporting that alterations in *ANGPTL6* might be a genetic risk factor influencing IA development.^20,21^

ANGPTL6 is one of the eight members of the secreted glycoprotein ANGPTL family, which share a common structure consisting of an amino-terminal coiled-coil domain, a linker region and a carboxy-terminal fibrinogen-like domain.^22^ In human, ANGPTL6 is described as a circulating protein mainly secreted from the liver and acting as an endocrine factor in the peripheral tissues.^23^ ANGPTLs are described as regulators of glucose and lipoprotein metabolisms through their amino terminus, and of angiogenesis through their carboxyl terminus.^24^ However, a large-scale population study showed that ANGPTL6 does not affect lipoprotein profile.^25^ In transgenic mice, expression of ANGPTL6 in epidermal keratinocytes promotes new blood vessel formation in the skin and vascular hyperpermeability.^22^ Homozygous *Angptl6* knockout mice die around E13 with cardiovascular defects and survivors develop obesity, hyperglycemia and hyperinsulinemia, and insulin resistance.^23^

Among ANGPTL6 variants identified in IA-affected family, the c.1378A>T nonsense mutation in the last exon of *ANGPTL6*, encodes a truncated ANGPTL6 protein lacking the last 11 C-terminal.^14^ We showed a 50% reduction of ANGPTL6 serum concentration in individuals heterozygous for the c.1378A>T allele compared to their non-carrier relatives. We demonstrated that this reduction results from the inability of the truncated protein lacking the last 11 amino acids (K460*-ANGPTL6) produced by the transcripts carrying the 1378A>T variant to be secreted.^14^ The 1378A >T *ANGPTL6* variant thus leads to ANGPTL6 haploinsufficiency, due to the inability of the truncated protein to be secreted.

The aim of this study is now to understand why the ANGPTL6 protein mutant predisposes to IA by combining analysis in *Angptl6*-knock in (KI) mice carrying the variant with *in vitro* studies to analyze the role of ANGPTL6 in vascular cells.

## Methods

All data needed to support the conclusions of this study are present in the paper and/or the Supplementary Material or are available via the corresponding authors upon reasonable request.

### Animal model

*Angptl6*-knock-in (KI) mouse model was generated by homologous recombination, by the Institut Clinique de la Souris (Illkirch, France), on a genetic background C57BL/6J. Thanks to the sequence homology between mice and human ANGPTL6, the p.Lys447Ter-ANGPTL6 point mutation introduced in mice mimics the human rare variant the p.Lys460Ter-ANGPTL6 predisposing to IA,^14^ both leading to a deletion of the last 11 amino acids. We generated control (*Angptl6^WT/WT^*), heterozygous (*Angptl6^WT/Δ^*), and homozygous (*Angptl6^Δ/ Δ^*) mice. The mice were housed under 12 h light/12 h dark cycles at 25 °C with free access to food and water. All animal care and use procedures were performed in accordance with the European Community Standards on the Care and Use of Laboratory Animals (Directive 2010/63/EU) and were approved by the local ethics committee (Comité d’Ethique en Expérimentation Animale des Pays de Loire: CEEA-006; authorization number: APAFIS#23002-2019112518261112 v2 and 00909.01).

### Hypertension model

The N-Nitro-L-Arginine methyl ester hydrochloride (L-NAME, Sigma) was used at a dose of 3 g/L in the drinking water for 21 days. Systolic blood pressure was monitored by non-invasive method (BP-2000 series II - Visitech systems)^26^ 2 to 3 times a week. All the mice on the hypertensive diet were measured for a fortnight under basal conditions and then 21 days under treatment.

### Aneurysm model

Mice between 8 and 10 weeks of age were subjected to ligations of the left common carotid arteries and posterior branches of right renal arteries under general anesthesia with 1% to 2% isoflurane. We took special care to maintain normal body temperature and not to induce excessively deep anesthesia on operation. One week after the first operation, the posterior branches of the left renal arteries were ligated. One week after these procedures, 0.9% saline was substituted for drinking water.^27^ All surviving mice were killed 4 months after the surgical procedure. In both groups, systolic blood pressure was measured by the tail-cuff plethysmography method before operation, after 2 months and before sacrifice.

### Brain collection and micro-computed tomography (micro-CT)

After deep anaesthesia with ketamine, mice were perfused with heparinized phosphate-buffered saline for 5 minutes and subsequently with 4% paraformaldehyde in phosphate buffer for 10 minutes and with a mix of gelatin (G1890; Sigma, 3% final) and barium sulfate (Micropaque®, Guerbet) for micro-CT observation or with gelatin-blue Evans solution. The brains were isolated and immersed in the same buffer for fixation during 24 h. Micro-CT scans were performed at the SC3M facility (Nantes, France) using a Skyscan 1272 (Bruker, Germany) microtomodensitometer at 12 µm resolution, 80kV (filter Al 1mm,) or 90kV (filter Al 0.5 mm + Cu 0.038 mm), rotation step: 0.35°, binning: 4. Three-dimensional reconstructions of the raw projections were obtained using Nrecon (Bruker) Smoothing: 0; Rig Artfact: 5; Beam Hardening: 10%. Using dedicated software (CTvox, CTan, Data Viewer from Bruker), internal diameters of cerebral arteries were measured.

### Pressure arteriography

Mice were sacrificed and their brains were harvested. Arterial segments of the posterior cerebral artery were dissected, cannulated on two glass micropipettes in an organ chamber containing physiological salt solution (PSS) maintained at 37 °C (pH 7.4), and pressurized using an arteriograph system (Living Systems Instrumentation, Inc., St. Albans, VT) as previously described.^28^ Once prepared, arteries were allowed to stabilize for at least 60 min at a pressure of 50 mmHg until the development of basal tone. Artery viability was tested using a KCl-rich solution (80 mmol/L) and PhE; 10 μmol/L). Flow was increased from 3 to 10 µl/min. Myogenic tone was determined by increasing intraluminal pressure by steps from 20 to 100 mmHg. Vessel internal diameter was continuously recorded using a CCD camera and edge-detection software. Diameters measured in PSS were considered active diameters. At the end of each experiment, maximal dilation was obtained in nominally Ca^2+^-free PSS containing EGTA (2-5 mmol/L; Sigma). Artery diameters are given in µm. Myogenic tone was expressed as the percentage of passive diameter ([passive diameter-active diameter]/passive diameter).

### Immunofluorescence staining

Retinas were collected on 6-day postnatal (P6) euthanized mice and fixed with 4% PFA in PBS overnight at 4°C for immunofluorescent staining. Blocking and permeabilization were performed using blocking buffer consisting of 1% FBS, 3% BSA (Sigma-Aldrich), 0.5% Triton X-100 (Sigma-Aldrich), 0.01% sodium deoxycholate (Sigma-Aldrich), and 0.02% sodium azide (Sigma-Aldrich) in PBS at pH 7.4 for 1 h at room temperature. DAPI (Sigma-Aldrich, 1/10000) was used for nuclear labelling and Isolectin B4 488 (life technologies, 1/400) was used to label endothelial cells. Retinas were mounted in Mowiol.

For labelling of vibratome sections, the right and the left parts of the brain were embedded separately in low melting point agarose (4%; Sigma-Aldrich). Each side was cut into 100 μm thick slices using a vibratome (Leica VT1200S, Leica Biosystems®). Slices were then permeabilized for 1 h with blocking buffer. Primary antibodies, Rabbit anti-SM22α (14106; Abcam, 1/1000) and Goat anti-VE-Cadherin antibody (AF938; RnD, 1/100), were incubated in 1:1 blocking buffer/PBS. Secondary antibodies, Donkey anti-rabbit alexa 568 and Donkey anti-goat alexa 488 (Invitrogen, 1/1000), were incubated in 1:1 blocking buffer/PBS. DAPI (1/1000) was used for nuclear labelling. Slices were put in µ-Slide 8 Well (80826; IBIDI®), with a drop of PBS on top of it before imaging.

### Brain transparization

Brains were render transparent using the CLARITY protocol. Briefly, electrophoresis (3h, 1.2A, 37°C, 100rpm) was applied, tissue was then permeabilized for 24h with blocking buffer (RT along the day and 4°C at night). Primary antibodies, Rabbit anti-SM22α (14106; Abcam, 1/200) was incubated in 1:1 blocking buffer/PBS with rocking at 4°C for 3 days and secondary antibodies, Donkey anti-rabbit alexa 568 was incubated at 1/400 concentration both in 1:1 blocking buffer/PBS for 2 days at 4°C. Isolectin B4 488 (life technologies, 1/400, 2 days) was used to label endothelial cells. Brains were placed back X-Clarity mounting solution before imaging.

### Fluorescent imaging

Images from fluorescently labelled arteries and retinas were acquired using a confocal A1 SIM microscope (Nikon) equipped with a Plan-Apochromat 20×/0.8 NA Ph2 objective or with a 60x/1.4 plan apochromat objective; or using an Eclipse Ti2 inverted microscope (Nikon). The microscopes were equipped with a photon multiplier tube detector. Images were taken at room temperature using NIS element software (Nikon).

### Light sheet microscopy

Imaging of transparized brains was performed in X-Clarity mounting solution to visualize the arteries of the circle of Willis and specifically the anterior-olfactory and anterior-middle cerebral artery bifurcations. The data were acquired on a ZEISS Lightsheet 7 with a Clr Plan-Neofluar 20×. The data were processed with ZEN imaging software and FIJI.

### Electron microscopy

Mice were deeply anesthetized and transcardially perfused with 15 ml of 0.1 M sodium phosphate buffer (PBS) containing 10 IU/ml heparin followed by 20 ml of 2.5% glutaraldehyde/4% paraformaldehyde in PBS 0.1M pH7.4. Mice were then perfused with mix gelatin/Evans blue (3%) to enable visualization the cerebral arteries. The skull was opened and incubated in 2% glutaraldehyde/2% paraformaldehyde in PBS for fixation of the brain tissues (1 h at room temperature then overnight at 4°C). The cerebral arteries were then dissected under a microscope (middle cerebral arery, anterior cerebral artery and olfactory artery) and post-fixed by incubation in 2% osmium tetroxide in PBS. Arterial samples were dehydrated by passage through a graded series of ethanol concentrations and were then embedded in epoxy resin. Semi-thin and ultrafin-thin sections were cut with an ultramicrotome (UC7, Leica), stained with 1% toluidine blue and screened by light microscopy to select areas of interest. Ultrathin sections of regions of interest were cut, mounted on copper grids, contrast stained with uranyl acetate and lead citrate and examined by transmission electron microscopy (JEM 1400, JEOL).

The average thickness of the media and intima were calculated using the area covered by the cells (smooth muscle or endothelial cells, respectively) over the total length of the bifurcation and for linear parts, over a 150 µm length of linear portion of the artery at least 200 µm away from the bifurcation.

### Cell culture

Primary rat aortic vascular smooth muscle cells (VSMCs) were isolated from aorta of Wistar Han rats. Explants were cleaned and digested for 2 h with collagenase II (1 mg/mL, Worthington Bio-chemical) at 37°C under agitation. Cells were cultured in Dulbecco modified Eagle medium (DMEM, Gibco; Invitrogen) containing 10% fetal bovine serum (FBS), 4.5g/L glucose, 100 units/mL penicillin and 100 μg/mL streptomycin at 37°C and 5% CO2. All experiments were performed between passages 2 and 4. Human aortic smooth muscle cells (C-12532, PromoCell) were cultured with Smooth Muscle Cell Growth Medium 2 (C-22062, PromoCell).

### Cell adhesion assay using impedance measurement

Plates were coated with 0.2% gelatine with or without recombinant ANGPTL6 protein (LS-G16246; LSBio) before experiment. VSMCs were seeded in a 96 well plate microtiter xCELLigence assay plate (E-Plate, ACEA Biosciences Inc.) and placed on the Real-time xCELLigence Cell Analyzer (Roche Applied Science) platform at 37°C to measure the “cell index” (change in impedance compared to the reference point) every 5 min for a period of 30 h. Adhesion speed was measured as cell index slope.

### Transwell migration assay

SMCs were added to the upper chamber of a Boyden chamber, (polycarbonate membrane, 8 µm pore size and 0.47cm^2^ of culture area (140629, Nunc™)) coated with 0.2% gelatin with or without ANGPTL6 recombinant protein at 5 ng/mL. The chemoattractant used was 10% FBS. Twenty-two hours after seeding, cells were fixed with 4% paraformaldehyde and stained with Coomassie blue 0.1%. Cells that stayed in the upper chamber were scrapped away and cells on lower side were observed using brightfield microscopy. Quantification of area covered by cells was done with FIJI.

For transmigration assay using impedance measurement, VSMC were seeded in the upper chambers of 16 well CIM-plates (ACEA Biosciences Inc.) coated with 0.2% gelatin with or without recombinant ANGPTL6 protein (LS-G16246; LSBio, for rat VSMC and AG-45A-0016YEK-KI01; Adipogen, for human VSMC) before experiment. VSMCs were left adhere to the coating in the incubator for 30 min. The plate was then placed on the Real-time xCELLigence Cell Analyzer (Roche Applied Science) platform at 37°C to measure the “cell index” (change in impedance compared to the reference point) every 5 min for a period of 15 h. The rate of migration was measured as cell index slope.

### Surface plasmon resonance (SPR) studies

SPR was used to assess the interaction between fibronectin (ECM001, Sigma-aldrich),

αVβ5 integrin (2528-AV, R&D systems) and α5β1 integrin (3230-A5, R&D systems) with ANGPTL6 (CSB-MP822824HU, Cusabio) immobilized onto CM5 (1000 RU; GE Healthcare) using a Biacore T200 (GE Healthcare). Single-cycle kinetics was performed by 2 min-sequential injections at a flow rate of 40 µL/min of five increasing concentrations of αVβ5 or α5β1 integrins (12.5 nM; 25 nM; 50 nM; 100nM and 200 nM) or fibronectin (62.5 nM; 125 nM; 250 nM; 500 nM and 1000 nM) over the ANGPTL6-functionalized substrate.

Each sensorgram was corrected by subtracting a sensorgram obtained from a reference flow cell with no immobilized protein. Global fitting of the αVβ5 integrin data to a Langmuir 1:1 model was used to determine the association (*ka*), dissociation (*kd*) and the affinity constant (*K*D) with T200 Biaevaluation Software (Biacore). *K*D values for α5β1 and fibronectin were determined using the steady-state method.

### Western blot

VSMC were incubated on ice with lysis buffer supplemented with protease and phosphatase inhibitor cocktails (Sigma Aldrich) and sodium orthovanadate. Lysates were subjected to SDS-PAGE, transferred to nitrocellulose membranes, and incubated with specific antibodies: P-MYPT (1/1000; #3040, Cell Signaling Technology) and MYPT (1/1000; #2634, Cell Signaling Technology), SM22α (1/1000; #14106, abcam). Equal loading was checked by reprobing of the membrane with an anti-tubulin antibody (T9026, Sigma). Immune complexes were detected with appropriate secondary antibodies and enhanced chemiluminescence reagent (Clarity ECL BioRad). Protein band intensities were quantified using FIJI.

### RT-PCR

Total RNA was purified from cells using RNA plus kit (Macherey Nagel) according to manufacturer instructions. RNA (500 ng) was reverse-transcribed with M-MLV enzyme (28025021, Thermo Fisher Scientific). Real-time qPCR was performed on a 7900HT Fast Real-Time PCR System (Applied Biosystems) using SYBR Green Master Mix (4367659, Applied Biosystems) and primers listed in Table S2. Each sample was analyzed in triplicate. GAPDH was used as the reference gene and results are expressed according to the 2^-ΔΔCt^ method.

### Collagen gel contraction assay

VSMC were mixed with collagen gel working solution (CBA-201, Cell Biolabs) with or without ANGPTL6 recombinant protein at 10 ng/mL. The cell-collagen mixture was added into a 24-well plate and incubated at 37°C for 1h to allow collagen polymerization. Culture medium without 10 % FBS was added to the top of the collagen gel lattice. Two days after seeding, cells were treated or not with SVF (10%) or BDM (from CBA-201 kit), and the collagen gels were then released using a sterile spatula. Changes in the collagen gel size were measured after 48 h.

### Statistical analysis

Statistical analysis was performed using GraphPad Prism software. Details of the statistical test used for each experiment can be found in the figure legends.

## Results

### *Angptl6*-KI mice show dilated cerebral arteries and defect in adaptation to high blood pressure

*Angptl6*-KI mice developed normally and survived to adulthood without showing neither gross abnormalities, nor obvious behavioral or physical phenotype. In particular, body weight (Figure S1) as well as the plasma concentration of cholesterol and triglycerides (Figure S2) are similar between *Angptl6^WT/Δ^* and *Angptl6^Δ/D^* and *Angptl6^WT/WT^*mice. To characterize the cerebral vasculature of *Angptl6*-KI mice, we first analyzed by micro-CT the circle of Willis under basal condition. We did not detect any major anatomical changes, including the angles of arterial bifurcations (Figure S3), but we observed that arteries of *Angptl6*-KI mice have significantly larger diameters than control mice (Figure 1A).

**Figure 1.**
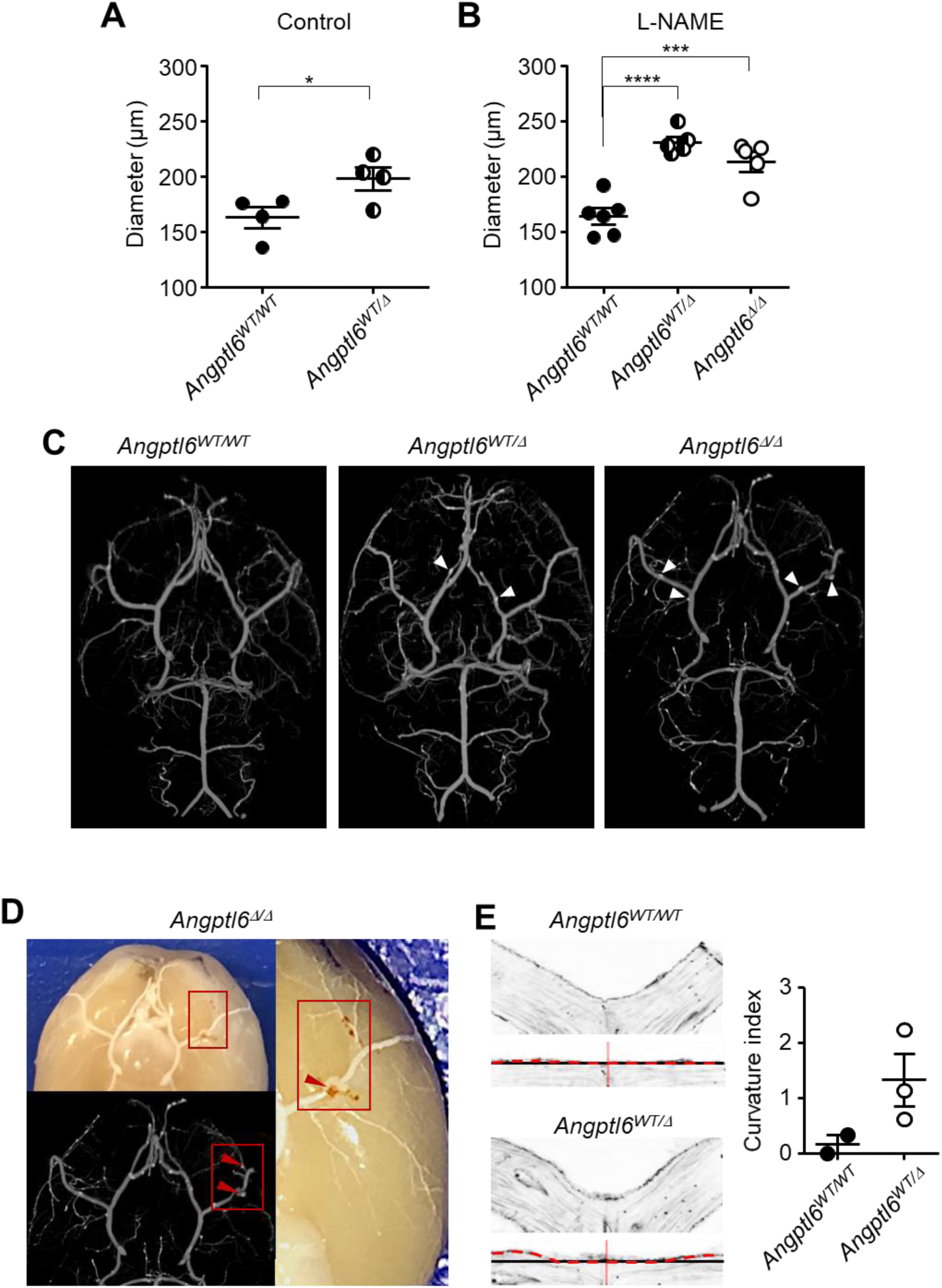
Basilar artery diameter under normotensive and hypertensive conditions in control and mutant mice. **A** Diameter of basilar artery of normotensive *Angptl6^WT/WT^*and *Angptl6^WT/Δ^* mice (6-8 months old). **B.** Diameter of basilar artery of normotensive *Angptl6^WT/^*^WT^, *Angptl6^WT/Δ^* and *Angptl6^Δ/Δ^* mice made hypertensive by treatment with L-NAME (3 g/L for 3weeks). Diameters have been measured on micro-CT images (one-way ANOVA; \**P*<0.05; \*\*\**P*< 0.001, \**P*<0.0001 vs. *Angptl6^WT/WT^* mice in the same condition; # *P*<0.05 vs. hypertensive *Angptl6^WT/Δ^* mice) **C.** Micro-CT images of the circle of Willis of *Angptl6^WT/^*^WT^, *Angptl6^WT/Δ^* and *Angptl6^Δ/Δ^* mice after 3 weeks of L-NAME treatment. White arrowheads indicate ectasia. **D.** Photograph and micro-CT of the brain of an *Angptl6^Δ/Δ^* mouse after 3 weeks of L-NAME treatment showing the colocalization of ectasia (red arrowheads) with blood hemorrhage. **E.** Images from vibratome sections of bifurcations of the circle of Willis of hypertensive *Angptl6^WT/WT^*and *Angptl6^WT/Δ^* mice (10-12 months old, 3 weeks of L-NAME treatment). For both genotype, below the native images of the bifurcation, same images are shown after digital linearization of the bifurcations. The apex of the bifurcation is indicated by the vertical red line. The blackline shows the theoretical straight line (Lt) and the dotted line followed the true delimitation of the bifurcation (Lex). Quantification of the local deformation was determined by the calculation of a curvature index as (Lex-Lt)/Lt) (*n*=3).

Because we previously found a significant difference in the rate of IA among individuals heterozygous for ANGPTL6 variants with a history of high blood pressure compared to normotensive ones, we next challenged the mice by inducing a mild hypertension.^14^ Administration of L-NAME for 3 weeks induced a similar rise of systolic blood pressure (∼15 mm Hg) in the three groups of mice (Figure S4). This rise in pressure did not modify the basilar artery diameter in in *Angptl6^WT/WT^* mice while it induced a significant increase in the diameter of basilar artery of *Angptl6^WT/Δ^* and *Angptl6^Δ/D^* mice, thereby majoring the difference in the arterial caliber between control and mutant mice (Figure 1B). In addition, examination of micro-CT images revealed the presence of small local dilations (*i.e.* ectasias) along arteries of the circle of Willis of *Angptl6^WT/Δ^* and *Angptl6^Δ/D^* mice which were never observed in *Angptl6^WT/WT^* mice (Figure 1C). Some of these ectasias were found to be co-localized with local blood leakage observed on the surface of the brain (Figure 1D). A fine examination of arterial bifurcations of the circle of Willis also revealed specific dilation of the arterial wall close to the apex in mutant mice, at sites where IA are known to develop (Figure 1E). These results thus show that, under basal condition, arteries of the circle of Willis of *Angptl6*-KI mice are inherently larger in diameter than those of control mice and that, unlike the latter, they further dilated in high blood pressure condition, suggesting a defect in their ability to adapt to pressure.

### ANGPTL6 variant promotes endothelial dysfunction in cerebral arteries

We then used pressure arteriography to analyze reactivity and passive properties of perfused posterior cerebral arteries *ex-vivo* (Figure 2). Increasing the perfusion pressure from 20 to 100 mmHg in the absence of external calcium caused a gradual increase in the passive artery diameter of both *Angptl6^WT/WT^* and *Angptl6^Δ/Δ^* mice. However, for all pressure values, arteries from *Angptl6^Δ/Δ^* mice were significantly larger than those from *Angptl6^WT/WT^* mice artery (Figure 2A). As illustrated for the measurements made at 50 mm Hg, this increase in passive diameter was similarly observed in arteries of *Angptl6^WT/Δ^* and *Angptl6^Δ/Δ^* compared to *Angptl6^WT/WT^*mice (Figure 2B). In contrast, we did not observe any difference in the myogenic response in the 3 groups of mice as well as in the isometric contraction induced by KCl and vasoconstricting agonists (PhE/5-HT) (Figure 2C). We next analyzed the flow-mediated dilation (FMD) in control and mutant mice. Stepwise increases in intraluminal flow in perfused posterior cerebral arteries induced vasodilation (Figure 2D). FMD was significantly reduced in cerebral arteries isolated from *Angptl6^Δ/Δ^* and *Angptl6^WT/Δ^* mice compared to *Angptl6^WT/WT^*mice (Figure 2D). To further characterized this change in FMD associated with Angptl6 mutation, FMD was measured in the presence of L-NAME to abolish its NO-dependent component. As expected, FMD was reduced in the presence of L-NAME, but the FMD curves in arteries from *Angptl6^WT/WT^*, *Angptl6^WT/Δ^* and *Angptl6^Δ/Δ^* mice were superimposed (Figure 2E), indicating similar remaining NO-independent component of FMD in mutant as in control mice. The reduced FMD in ANGPTL6 mutant mice was thus due to a decrease in the NO-dependent FMD (Figure 2F). Taken together, the observed increase in the passive internal diameter of *ex-vivo* cerebral arteries of mutant mice confirmed observations made in micro-CT images, suggesting that Angptl6 mutation induces intrinsic change in cerebral artery wall structure, associated with endothelial dysfunction characterized by a decreased NO production in response to flow. Measurement of contraction and NO-dependent relaxation of extracerebral arteries (mesenteric), did not reveal any difference between *Angptl6*-KI and *Angptl6^WT/WT^* mice (Figure S5).

**Figure 2.**
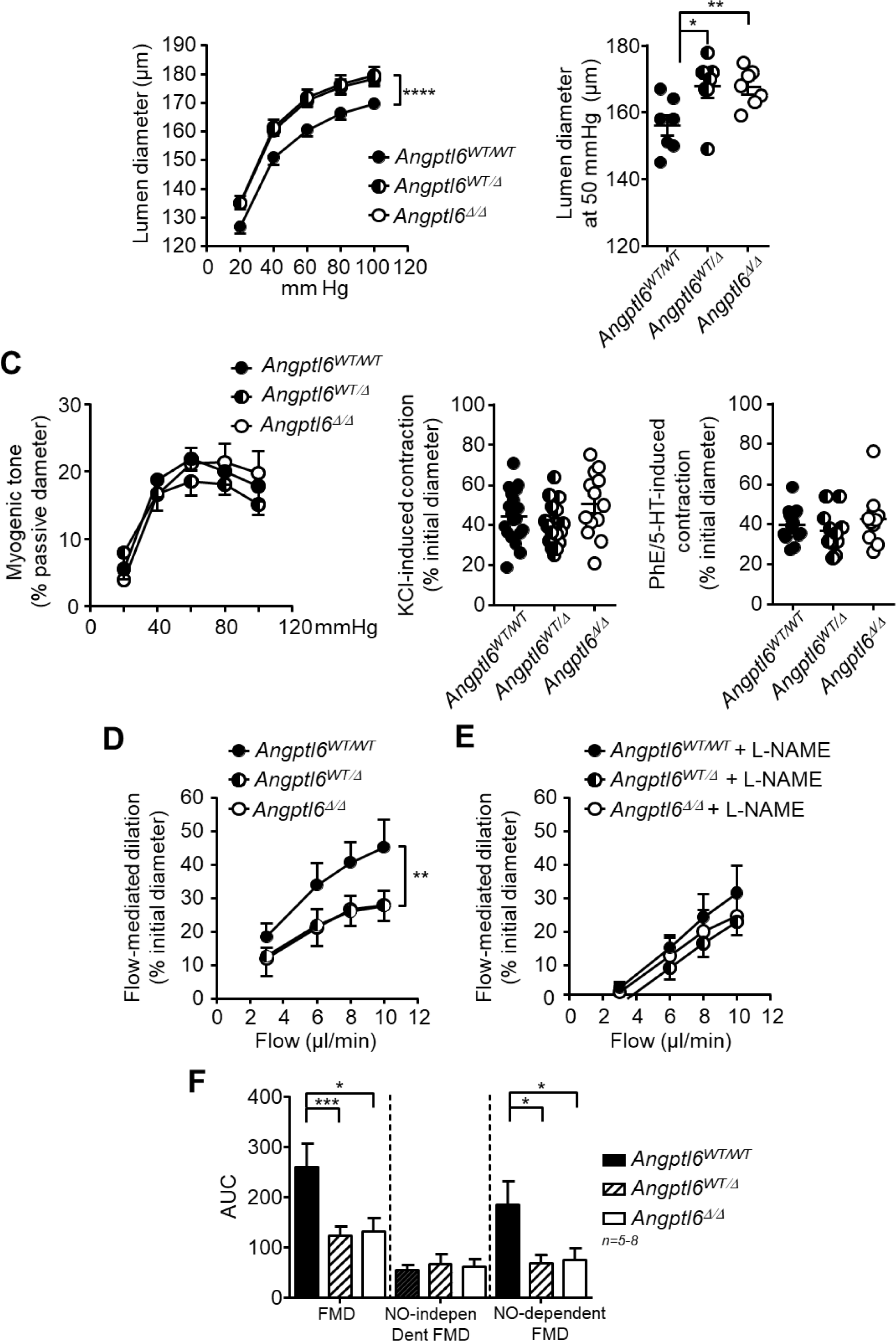
Endothelial dysfunction of cerebral arteries of ANGPTL6 KI mice. **A.** Passive diameter-experimental perfusion pressure relationships of posterior cerebral artery of *Angptl6^WT/WT^* and *Angptl6^Δ/Δ^* mice in the absence of external calcium (n=7, Two-way ANOVA; \*\*\**P*< 0.001). **B.** Passive inner diameter of *Angptl6^WT/WT^*, *Angptl6^WT/Δ^* and *Angptl6^Δ/Δ^* mice at intraluminal pressure of 50 mmHg (One-way ANOVA; \*\**P*< 0.01, \*\**P*<0.01). **C.** Pressure-induced myogenic tone in posterior cerebral artery of *Angptl6^WT/WT^*, *Angptl6^WT/Δ^* and *Angptl6^Δ/Δ^* mice expressed as % of passive diameter (n= 8-10 mice; 2-way ANOVA, ns between genotype) and contraction of posterior cerebral artery of *Angptl6^WT/WT^*, *Angptl6^WT/Δ^* and *Angptl6^Δ/Δ^* mice induced by KCl or vasocontracting mediator (PhE/5-HT) expresse as % of the initial diameter (One-way ANOVA; ns) D. Flow-mediated dilation (FMD) of posterior cerebral artery of *Angptl6^WT/WT^* and *Angptl6^Δ/Δ^* mice in the absence and in the presence of L-NAME (n=5). **E.** Quantification of the FMD by the area under the dilation-flow curves (AUC) under basal condition, in the presence of L-NAME (NO-independent FMD) and the subtraction of the AUC in the presence of L-NAME from the AUC under basal condition (NO-dependent component) in *Angptl6^WT/WT^* and *Angptl6^Δ/Δ^* mice. (n=5, Mann-Whitney test; \**P*<0.05; \*\**P*< 0.01).

### Effect of ANGPTL6 mutation on retinal angiogenesis

ANGPTL6 has been shown to promote angiogenesis.^22^ We therefore use the retinal vascular development in neonatal mice (P6) as a model to assess whether the *Angptl6* variant affected angiogenesis. Imaging of the vascular network by endothelial labelling did not reveal differences in the vascular density as well as the radial expansion of the retinal vascular network between *Angptl6^WT/WT^*, *Angptl6^Δ/Δ^* and *Angptl6^WT/Δ^* mice (Figure 3A and B). This suggests that the *Angptl6* variant did not interfere with sprouting, migration and proliferation of endothelial cells at the vascular front.

**Figure 3.**
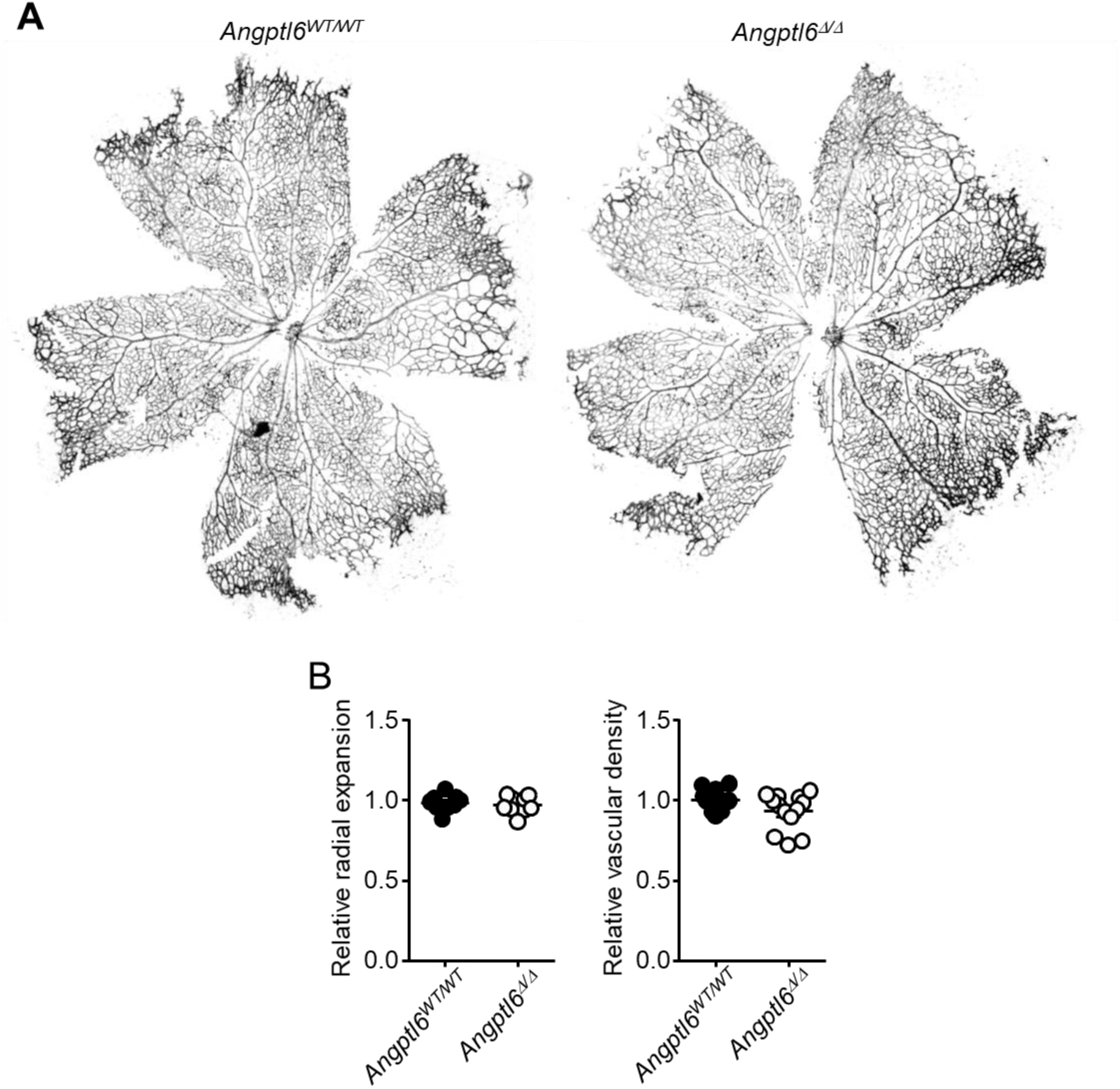
Retinal angiogenesis in *Angptl6^WT/WT^* and *Angptl6^WT/Δ^* neonates. **A.** Representative images of retinal vasculature after fluorescent labelling of endothelial cells (isolectin B4) of *Angptl6^WT/WT^, Angptl6 ^WT/Δ^* and *Angptl6^Δ/Δ^* mice (P6 neonates). **B.** Corresponding quantification of radial expansion and vascular density. (Mann-Whitney test; ns).

### Effect of ANGPTL6 mutation on IA formation

To assess whether ANGPTL6 mutation could potentiate IA formation in mice, we used the previously described model of IA induction through increasing hemodynamic stress at bifurcation sites of cerebral arteries.^27^ As previously described, examination of the circle of Willis 4 month after surgery shows typical enlargement and tortuosity of cerebral arteries (Figure 4A). Quantification of the anterior cerebral artery diameter clearly demonstrated the significant dilation of the artery induced by the hemodynamic overload (Figure 4B).

**Figure 4.**
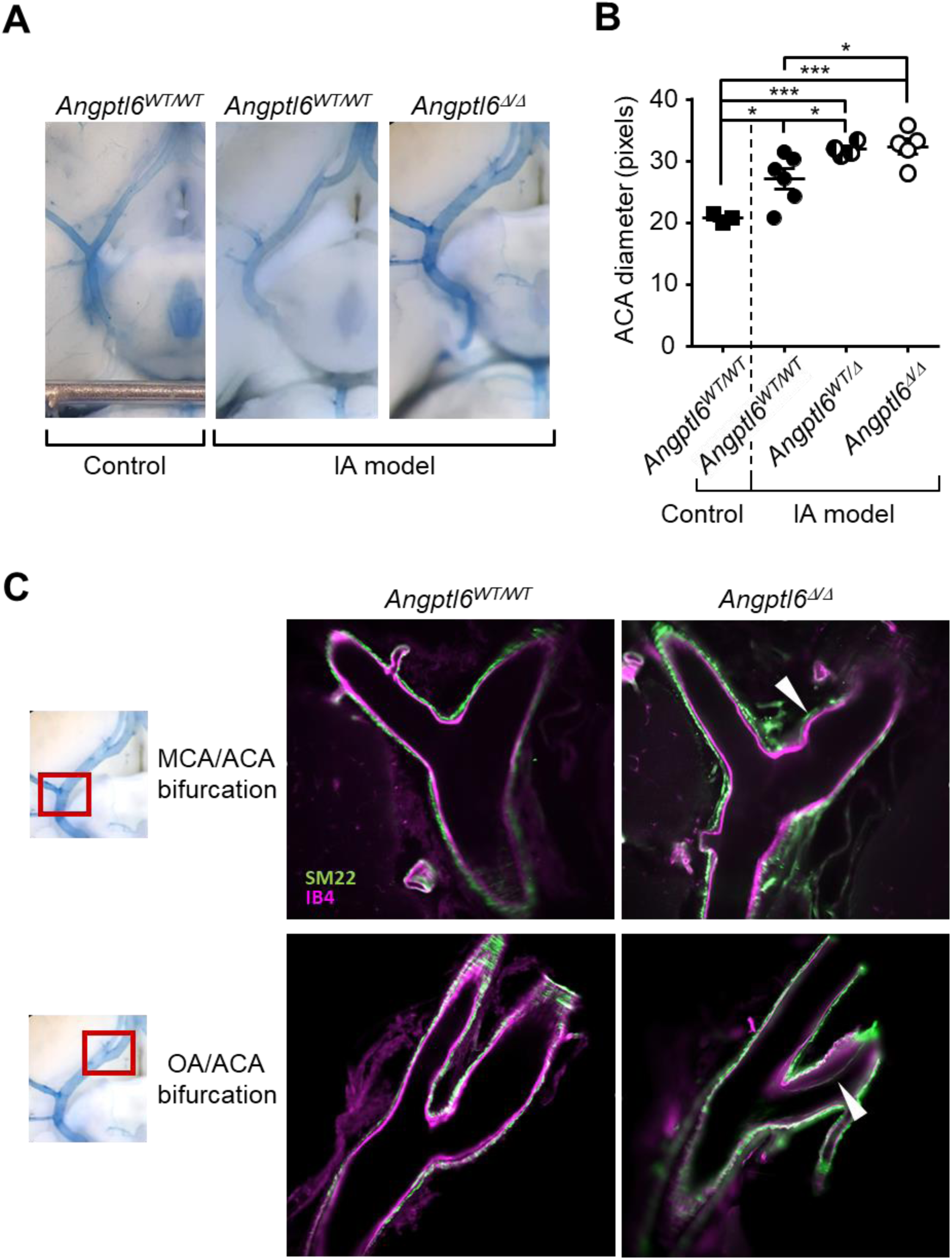
IA model in control and mutant mice. **A.** Representative gross images of cerebral artery in control condition (*Angptl6^WT/WT^* mice) and 4 months after the surgery to induce IA (*Angptl6^WT/WT^* and *Angptl6^Δ/Δ^* mice). **B**. Measurement of the diameter of the anterior cerebral artery in control condition (*Angptl6^WT/WT^* mice) and 4 months after the surgery to induce IA (*Angptl6^WT/WT^*, *Angptl6^WT/Δ^* and *Angptl6^Δ/Δ^* mice) (One-way ANOVA; \*\**P*< 0.01, \*\*\**P*< 0.001). **C**. Representative light sheet images of cerebral bifurcations of *Angptl6^WT/WT^*and *Angptl6^Δ/Δ^* 4 months after the surgery to induce IA, after fluorescent labelling of smooth muscle cells (SM22α, green) and endothelial cells (isolectin B4, Magenta).

Interestingly, this dilation was stronger in *Angptl6^WT/Δ^* and *Angptl6^Δ/Δ^* compared to *Angptl6^WT/WT^*mice (Figure 4A and B). A detailed assessment of arterial bifurcations by light sheet microscopy after labelling of endothelial cells and VSMC revealed structural changes only in *Angptl6*-KI mice, which were similar to those described at early stage of IA formation in humans. We observed typical enlargement and deformation of the wall close to the apex of the bifurcation as well as intimal hyperplasia, with disorganization of the media layer (Figure 4C).

To define more precisely the arterial wall alterations induced in the IA model among the 3 groups of mice, we analyzed linear segments and bifurcations of the anterior cerebral artery and the olfactory artery using electron microscopy (Figure 5). The linear portion of the artery wall in the control mice has a typical structure, with a thin and homogeneous intima, a continuous internal elastic lamina (IEL) and a media composed of well-organized, contiguous VSMC oriented perpendicular to the lumen axis (Figure 5A). At the bifurcation, this organization is altered, with typical breaks in the IEL and focal thickening of the intima adjacent to the bifurcation apex. This wall remodeling is strongly aggravated in homozygous *Angptl6*-KI mice which show greater cell proliferation and neointimal thickening, in addition to ruptures of the IEL, in the area adjacent to the bifurcation apex (Figure 5A). The media also showed remodeling manifested by thickening and disorganization of VSMC. Moreover, even the linear portions of the arterial wall show altered structure in Angptl6-KI mice, with an irregular, thickened intima, and a media composed of distorted and disorganized VSMC. Measurement of intimal and media thickness confirms these observations, with significantly greater thickening in homozygous Angptl6-KI mice than in control mice, with heterozygous Angptl6-KI mice showing intermediate behavior (Figure 5B). This aggravated arterial wall damage along the circle of Willis induced by hemodynamic overload in Angptl6-KI mice compared with control mice confirms previous observation in the context of high blood pressure (Figure 2A). These results thus suggest that *ANGPTL6* variants could favor IA formation by predisposing the arterial wall to deformations and intima and media remodeling at the bifurcations of the circle of Willis in condition of hemodynamic overload.

**Figure 5.**
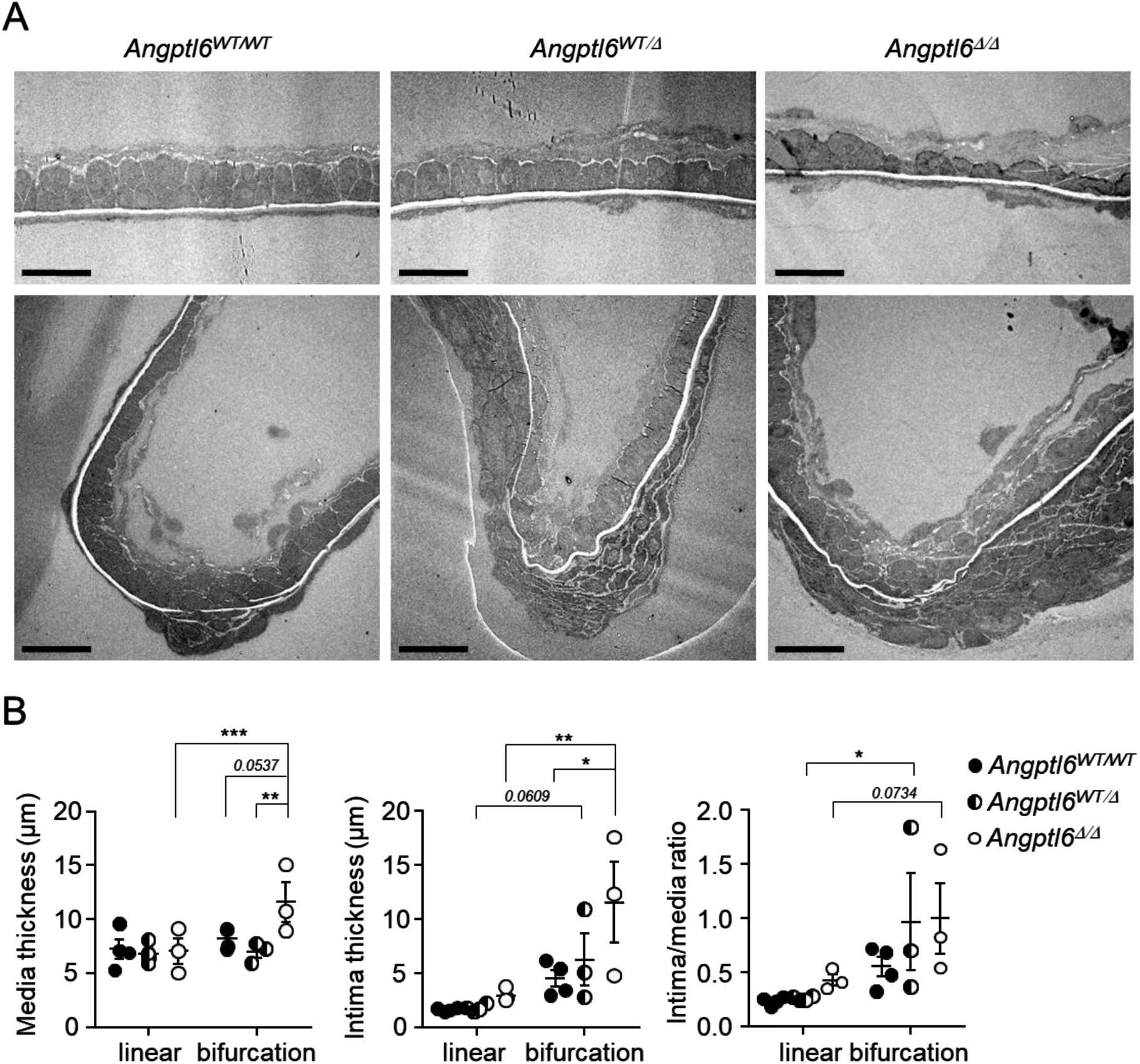
Electron micrographs of cerebral artery wall. **A.** Representative images of linear parts (top) and bifurcations (bottom) of the anterior cerebral artery and the olfactory artery 4 months after the surgery to induce IA in *Angptl6^WT/WT^*, *Angptl6^WT/Δ^* and *Angptl6^Δ/Δ^* mice (Bar: 10 µm). **B.** Quantification of media and intima thickness and intima/media ratio in the 3 groups of mice (Two-way ANOVA; \**P*< 0.05, \*\**P*< 0.01, \*\*\**P*< 0.001).

### ANGPTL6 regulates VSMC adhesion and migration

For further insight into the role ANGPTL6 in the media layer of the arterial wall, we directly analyzed its effect on VSMC. As ANGPTL6 is a secreted extracellular and circulating protein, we started by adding ANGPTL6 to the culture medium, but we were unable to detect any effect on VSMC functions (not shown). We then studied VSMC which were seeded on ANGPTL6-containing gelatin coating. Proliferation of VSMC was similar on gelatin alone or on ANGPTL6-containing gelatin (not shown). By contrast, ANGPTL6 coating accelerated the adhesion of VSMC, with an effect similar for concentrations ranging between 1 to 4 ng/mL (Figure 6A and B). A same effect of ANGPTL6-containing gelatin was observed in VSMC originating from human arteries (Figure S6). On the other hand, the presence of ANGPTL6 in the extracellular matrix inhibited VSMC migration (Figure 6C and D). ANGPTL6 thus appears as an extracellular matrix factor which increase VSMC adhesion and limits migration. This increased adhesion of VSMC induced by ANGPTL6 was further confirmed by a stronger activation of FAK signaling, monitored by FAK phosphorylation, in VSMC growing on ANGPTL6-containing matrix compared to VSMC seeded on gelatin alone (Figure 6E). This suggests that, in the other way, the absence of ANGPTL6 may lead to less adherent VSMC, more prone to migrate.

**Figure 6.**
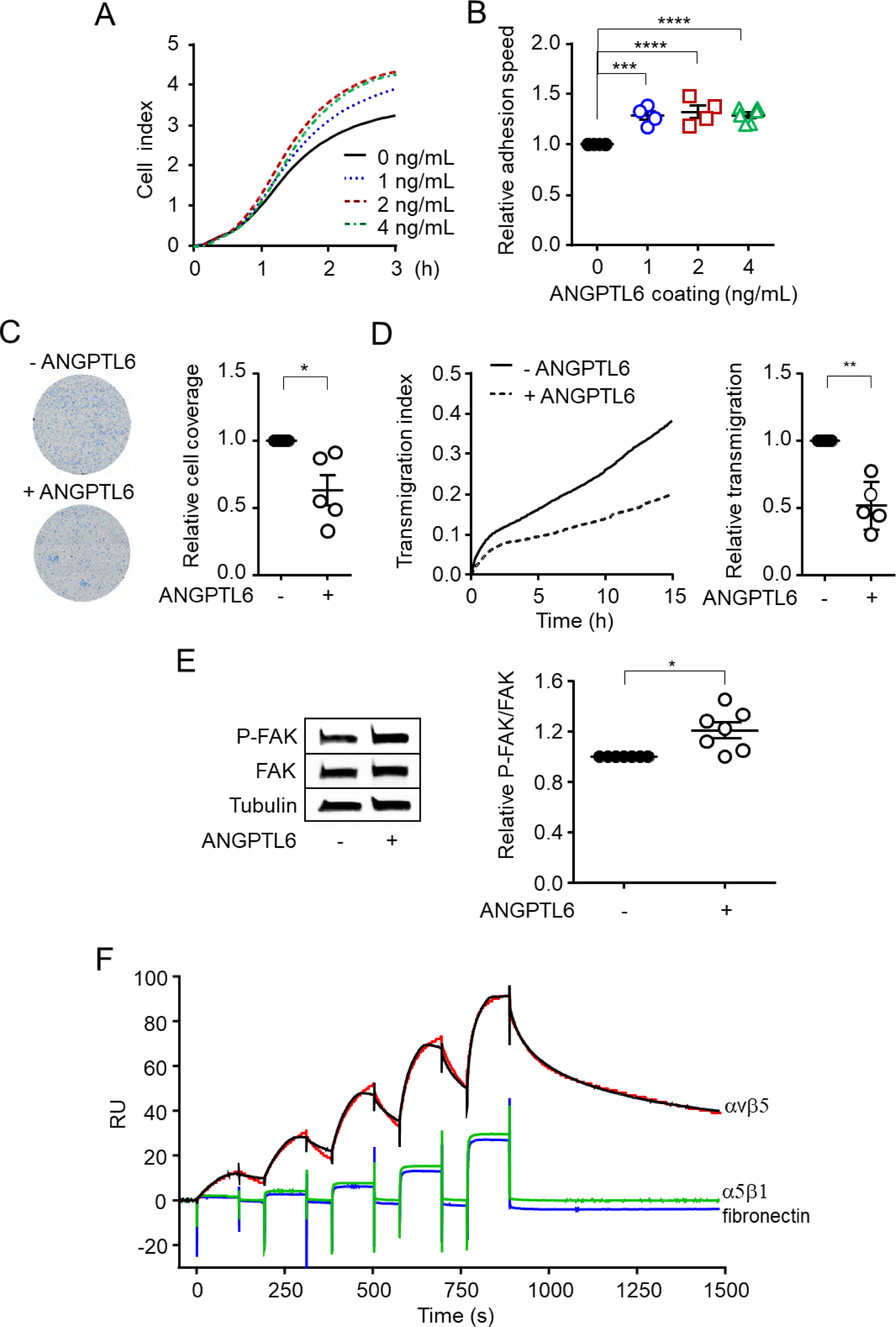
ANGPTL6 increases VSMC adhesion and reduces VSMC transmigration. **A.** Representative curve of cell index measurement for rat VSMC adhering on gelatin coating without (grey) and with ANGPTL6 (4 ng/mL, blue). **B**. Quantification of the rate of VSMC adhesion on gelatin coating without or with ANGPTL6 (1-4 ng/mL) using the slope of the cell index curve in its linear part (One-way ANOVA; \*\*\**P*< 0.001). **C.** Representative images of the evan blue staining of rat VSMC that transmigrated across a porous membrane coated with gelatin containing or not ANGPTL6 after 24 h (left) and the corresponding quantification of the area covered by rat VSMC (normalized to the gelatin alone condition, Wilcoxon *t*-test) **D.** Representative curve of cell index measurement for human VSMC adhering on gelatin coating without (grey) and with ANGPTL6 (4 ng/mL, blue) (left) and corresponding quantification of the rate of adhesion using the slope of the cell index curve in its linear part (Wilcoxon *t*-test; \**P*<0.05). **F.** Single-cycle kinetic SPR analysis of the ANGPTL6 interaction with αVβ5 or α5β1 integrins (12.5 nM; 25 nM; 50 nM; 100nM and 200 nM) or fibronectin (62.5 nM; 125 nM; 250 nM; 500 nM and 1000 nM). The red line represents the kinetic global fit of the ANGPTL6-αVβ5 interaction using 1:1 Langmuir model.

### ANGPTL6 interacts with αvβ5 integrin

ANGPTL6 presents a RGD (Arg-Gly-Asp) motif in its amino-acid sequence. The increased adhesion and FAK signaling activation in VSMC growing on ANGPTL6-containing gelatin may thus result from an increase of RGD-binding integrin engagement. To test this hypothesis, we used SPR to analyze potential direct interaction of ANGPTL6 with the major VSMC RGD-binding integrins, αvβ5 and α5β1. Sensorgram of αvβ5 integrin clearly exhibit typical binding response (Figure 6F). The analysis of the association of ANGPTL6 with αvβ5 integrin by using a global 1:1 Langmuir model of interaction revealed an equilibrium dissociation constant (*K*D) value in the nanomolar range (13 nM) (Fig. 6F). The reversible binding of ANGPTL6 with αvβ5 integrin is characterized by rapid association (association rate constant [*k*a] = 9.15 × 10^4^ m^−1^·s^−1^) and dissociation process (dissociation rate constant [*k*d] = 1.20 × 10^−3^·s^−1^). In contrast, sensorgrams of fibronectin and α5β1 integrin display almost square profiles with only slight curvature in the association phase consistent with a non-specific, low-affinity direct binding to ANGPTL6 (*K*D = 2 µM and 4 µM, for fibronectin and α5β1 integrin respectively; Fig. 6F). These data show that ANGPTL6 specifically and directly interact with αvβ5 integrin.

### ANGPTL6 does not affect VSMC phenotype but increases VSMC contractility

VSMC are capable of a remarkable plasticity, and phenotypic modulation of VSMC plays a significant role in IA formation and rupture.^29^ We thus assessed the potential role of ANGPTL6 on VSMC phenotype by measuring the expression of contractile and secretory phenotype marker gene expression (Figure 7A and B). The expression of all tested VSMC marker genes, but also genes related to extracellular matrix, was similar in VSMC cultured on gelatin and on ANGPTL6-containing gelatin. The equal expression level of SM22α protein in VSMC on gelatin and on ANGPTL6-containing gelatin confirm that ANGPTL6 had no effect on VSMC phenotype (Figure 7C). However, the level of phosphorylation of MYPT was increased in VSMC growing on ANGPTL6-containing matrix suggesting that ANGPTL6 stimulated the RhoA-Rho kinase-dependent contractility (Figure 7D). This was confirmed by gel contraction assay showing that VSMC contracted more strongly collagen-containing ANGPTL6 gel than a collagen gel alone (Figure 7E). As this increase in tension and MYPT phosphorylation induced by matrix ANGPTL6 was not associated with a change in the contractile phenotype and, as a consequence, the intrinsic contractile properties of VSMC, it may reflect a stronger a cell/matrix interaction due to ANGPTL6-integrin interaction.

**Figure 7.**
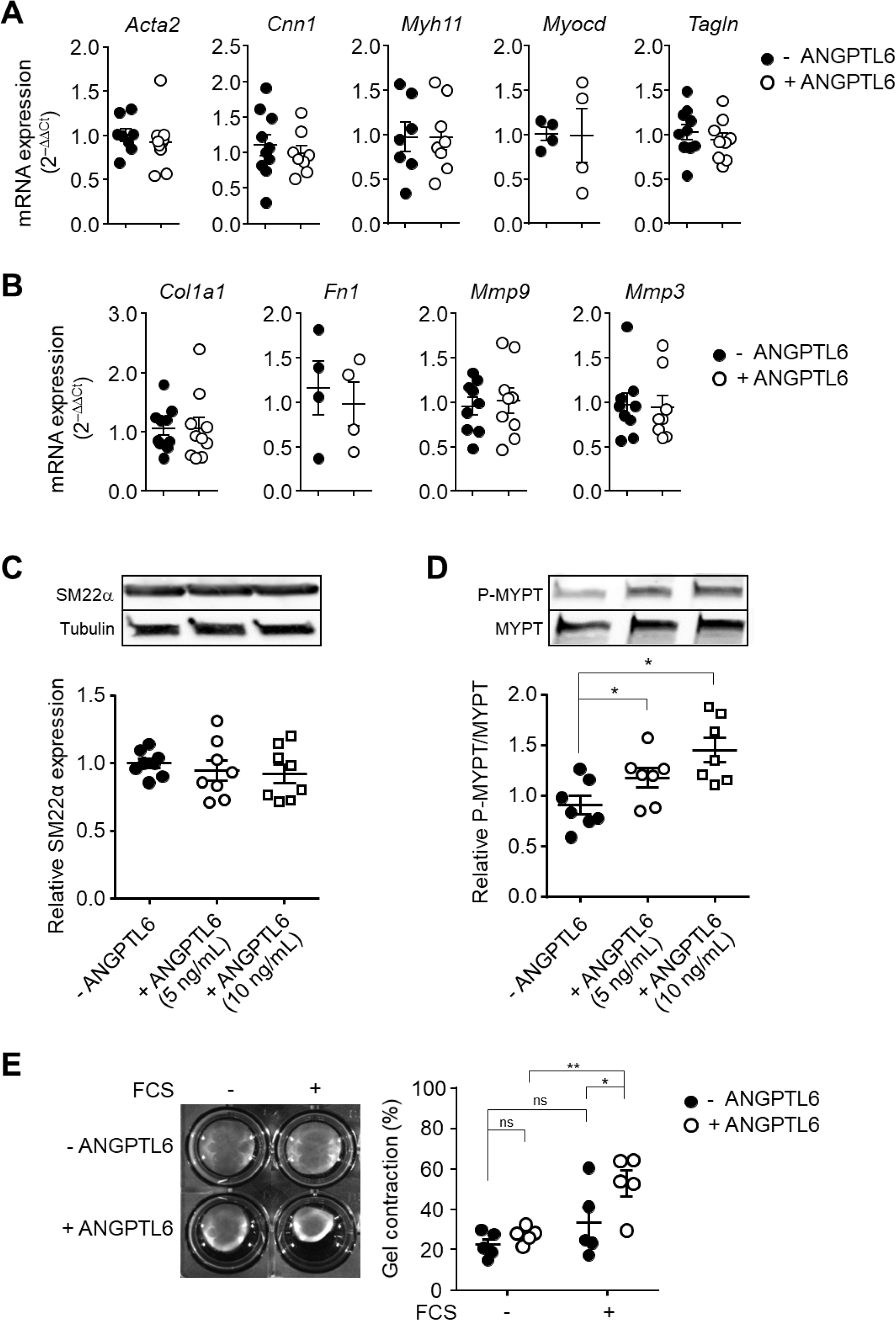
ANGPTL6 does not modulate VSMC phenotype but increases contractility. **A.** RT-qPCR analysis of mRNA levels of contractile phenotype marker genes (left) and synthetic phenotype marker gene (right) in VSMC cultured on gelatin alone and on AGPLT6-containing gelatin (5 ng/mL) **B.** Representative western blot analysis showing the expression of the contractile marker SM22 in VSMC cultured on gelatin alone and on AGPLT6-containing gelatin and corresponding quantification by densitometric analysis of SM22α bands relative to tubulin and normalized to the gelatin alone condition. **C.** Representative western blot analysis showing the expression of MYPT and phosphorylated MYPT (P-MYPT) the in VSMC cultured on gelatin alone and on AGPLT6-containing gelatin and corresponding quantification by densitometric analysis bands P-MYPT relative to MYPT and normalized to the gelatin alone condition. **D.** Representative images of gel contraction induced by VSMC seeded in collagen gels without or with ANGPTL6 (5 ng/mL) 48 h after release with and without serum (left) and corresponding quantification (right).

## Discussion

Our present study shows that the expression in mice of *ANGPTL6* variant predisposing to IA in humans is sufficient to 1) induce endothelial dysfunction expressed by reduced NO production in response to flow in cerebral arteries, 2) major dilatation of cerebral arteries, which may be accompanied by hemorrhage under hypertensive stress, and 3) aggravate wall damage and aneurysmal remodeling of the intima and media at arterial bifurcations of the circle of Willis under hemodynamic overload. *Angptl6*-KI mice thus reproduced key hallmarks of IA in humans. These observations, combined with *in vitro* experiments, allow us to document the role of ANGPTL6 in arteries and to propose an explanation for the increase in IA susceptibility induced by *ANGPTL6* variant. Moreover, in contrast to obese and insulin-resistant *Angptl6* knock-out mice,^23^ *Angptl6*-KI mice did not display any defect in lipid metabolism or metabolic syndrome-related phenotype. Although surprising, this result is consistent with both the absence of any lipid metabolic disorders in individuals carrying *ANGPTL6* variants in IA families,^14^ and the results from a large-scale population study that showed that ANGPTL6 levels does not affect lipoprotein profile.^25^ We can thus exclude a potential indirect role of a modified lipid metabolism in the vascular and IA-prone phenotype of *Angptl6*-KI mice.

The damage and dysfunction of endothelial cells are proposed to be an initial event in the pathogenesis of IA in humans.^30,31^ The endothelial dysfunction and reduced flow-induced NO-mediated dilation observed in *Angptl6*-KI mice are in agreement with the previously described stimulatory role of ANGPTL6 in endothelial NO production through the activation of ERK1/2 pathway.^32^ NO production induced by endothelial muscarinic receptor stimulation (Ach), was not modified in *Angptl6*-KI mice, suggesting that the regulatory role of ANGPTL6 on NO synthesis depended on the upstream signaling pathway engaged. Experiments performed in genetically modified experimental mouse models over-expressing ANGPTL6 in keratinocytes or using recombinant ANGPTL6 in both *in vitro and in vivo* assays also showed that ANGPTL6 induced angiogenesis and arteriogenesis through ERK1/2-NO pathway.^22,32^ However, angiogenesis in the developing retina was not altered in *Angptl6*-KI mice suggesting that endogenous ANGPTL6 did not play a major role in developmental or physiological angiogenesis. This absence of angiogenesis defect in *Angptl6*-KI mice is in agreement with their normal reproduction and growth. The role of ANGPTL6 on endothelial cells *in vivo* therefore seems subtle and limited to regulating the stimulation of NO production by flow. Thus, ANGPTL6 non-secreted variants would specifically contribute to an alteration in the physiological response of endothelial cells to flow, in line with the theory that considers IA as an acquired lesion resulting from a defective response of the vascular wall to local hemodynamic stress.^33^ This is supported by the exaggerated endothelium remodeling observed at bifurcation of the circle of Willis in *Angptl6*-KI mice under hemodynamic overload.

Regarding VSMC, we described for the first time the role of ANGPTL6 in this vascular cell type. ANGPTL6 modulates VSCM functions when incorporated to extracellular matrix. Matrix bound ANGPTL6 decreased VSMC migration and increased adhesion and FAK signaling activation, but did not affect VSMC phenotype. ANGPTL proteins, including ANGPTL6, are characterized by a N-terminal coiled-coil domain involved in lipid metabolism and a C-terminal fibrinogen-like domain containing an RGD motif, mediating receptor binding and other effects of ANGPTL,^34^ making integrins as good receptor candidates to mediate the effect of ANGPTL6 on VSMC. ANGPTL6 has been shown to stimulate adhesion of keratinocytes, fibroblasts and endothelial cells and this effect was antagonized by exogenous synthetic RGD peptides, suggesting the involvement of ANGPTL6-integrin interaction.^35^ In cancer cells, ANGPTL6 mediated its effects by binding to α6 integrin/E-cadherin complex.^36^ Here we show that ANGPTL6 binds the αvβ5 RGD-binding integrin thus identifying a new ANGPTL6 receptor. Integrins connect extracellular matrix proteins to the actin cytoskeleton clustered at the focal adhesion complex thus playing a crucial role in sensing ECM composition and mechanical strain and transducing mechanical cues to intracellular signals that produce functional adaptive response of VSMC.^37^ Activation of αvβ5 integrin is likely to be responsible for the increase adhesion and tension on VSMC induced by matrix-bound ANGPTL6. ANGPTL6 thus appears to be a novel regulator of vascular mechanotransduction, which supports the role of its variants in increasing IA susceptibility, by compromising an efficient and adapted response to hemodynamic stresses.

Our results thus identify ANGPTL6 as a protective component of the extracellular matrix of the media layer of cerebral arteries against pathological remodeling induced by hemodynamic overload. The media is accessible to circulating protein such as ANGPTL6, which are convected out of the blood through the vascular wall.^38^ High blood pressure increases this unidirectional outward convection of circulating factors from the blood across the arterial wall.^38,39^ That means that hypertension would lead to an increase in the amount of ANGPTL6 within the arterial wall, thus enhancing smooth muscle αvβ5 integrin-extracellular matrix interaction, to ensure effective mechanosensing and enable vascular smooth muscle cells to detect and adequately respond to change in intraluminal pressure. This adaptive mechanism would be defective in individuals with low circulating ANGPTL6 concentration as described in families with IA carrying the non-secreted ANGPTL6 variant,^14^ and probably in our model of *Angptl6-*KI mice, making them more susceptible to IA under high blood pressure condition. Unfortunately, despite considerable effort and multiple attempts with numerous antibodies, it was not possible to measure ANGPTL6 concentration in the blood of *Angptl6*-KI mice to confirm that as in human, it was decreased. Anyway, the proposed mechanism helps explain the role of hypertension in IA, particularly in individuals carrying ANGPTL6 variant^14^ and supports the concept that mechanotransduction disorders at the level of vascular smooth muscle cells of cerebral arteries play a role in the genesis of IA.^40^

## Acknowledgements

We are grateful to the animal house facility (UTE) of Nantes Université for their expert services. We acknowledge the IBISA MicroPICell facility (Biogenouest), member of the national infrastructure France-Bioimaging supported by the French national research agency (ANR-10-INBS-04) and of UMS BioCore (Inserm US16 et UAR CNRS 3556). We thank Joelle Vezier and the imaging platform SC3M (SFR Francois Bonamy, Nantes Université) for of micro-CT imaging, and Mike Maillasson and the Impact platform (SFR François Bonamy, Nantes Université) for surface plasmon resonance studies. We also acknowledge Florence Manero and the platform SCIAM (SFR ICAT, University of Angers) for electron microscopy.

## Author contributions

CM, MF, M-AR, MR, VL and TQ performed experiments, collected the data and analysed the results. CL performed experiments. HD, RR and RB participated to data interpretation and reviewed the manuscript. ACV designed the study, analysed the results, and reviewed the manuscript. GL conceived and directed the study and wrote the manuscript.

