## Supplemental Tables and Figures for "ANGPTL6 variant induces cerebral vascular dysfunction and predisposes to intracranial aneurysm in mice"

| Target gene | Forward primer (5'-3' sequence) | Reverse primer (5'-3' sequence) |
|---|---|---|
| *Gapdh* | AACCCATCACCATCTTCCAG | CCAGTAGACTCCACGACATAC |
| *Acta2* | ACGCGAAGCTCGTTATAGAAG | GACCCTGAAGTATCCGATAGAAC |
| *Cnn1* | GATCCACTCTCTCAGCTCCT | CTTCCGCACACTTTAACCGA |
| *Myh11* | CTTTCCAGCTCCAGACTCAC | CGCCTCACATCTATGCCATT |
| *Myocd* | CTGAGCAGTTGGAATGGATCT | CAAGGTCAGAAACAGATCGGA |
| *Tagln* | GCTCCTCATCATACTTCTTCTCA | AACGCTACTCTCCTTCCAG |
| *MMP3* | CTGTGGAGGACTTGTAGACTG | CTATTCCTGGTTGCTGCTCAT |
| *MMP9* | GGAGGTCATAGGTCACGTAGG | GAACTCACACAACGTCTTTCAC |
| *Fn1* | CGAGGTGACAGAGACCACAA | CTGGAGTCAAGCCAGACACA |
| *Col1a* | TACAGCACGCTTGTGGATGG | CAGATTGGGATGGAGGGAGTT |

**Table S1 Sequence of qRT-PCR primers**

### Supplemental Figures

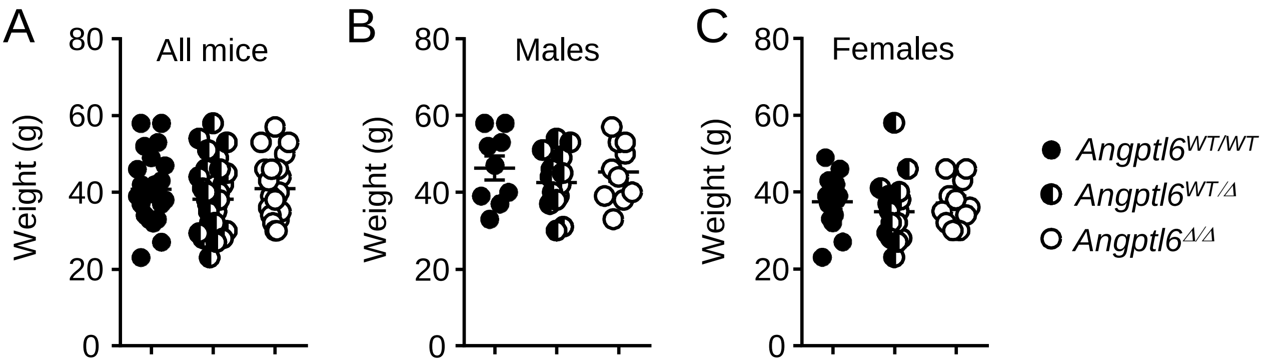


**Figure S1.** Body weight of *Angptl6*$^{WT/WT}$, *Angptl6*$^{WT/\Delta}$ and *Angptl6*$^{\Delta/\Delta}$ mice. **A**, All mice (mean age: 19.05±0.21 m, 19.30±0.37 m and 18.84±0.34 m, respectively). **B**, Male mice (mean age: 19.24±0.28 m, 19.97±0.21 and 19.72±0.30 m, respectively). **C**, Female mice (mean age: 18.93±0.31 m, 18.74±0.64 and 18.05±0.0.48 m, respectively). (*ns* between genotype and sex, One -way ANOVA, Bonferroni post-hoc test).

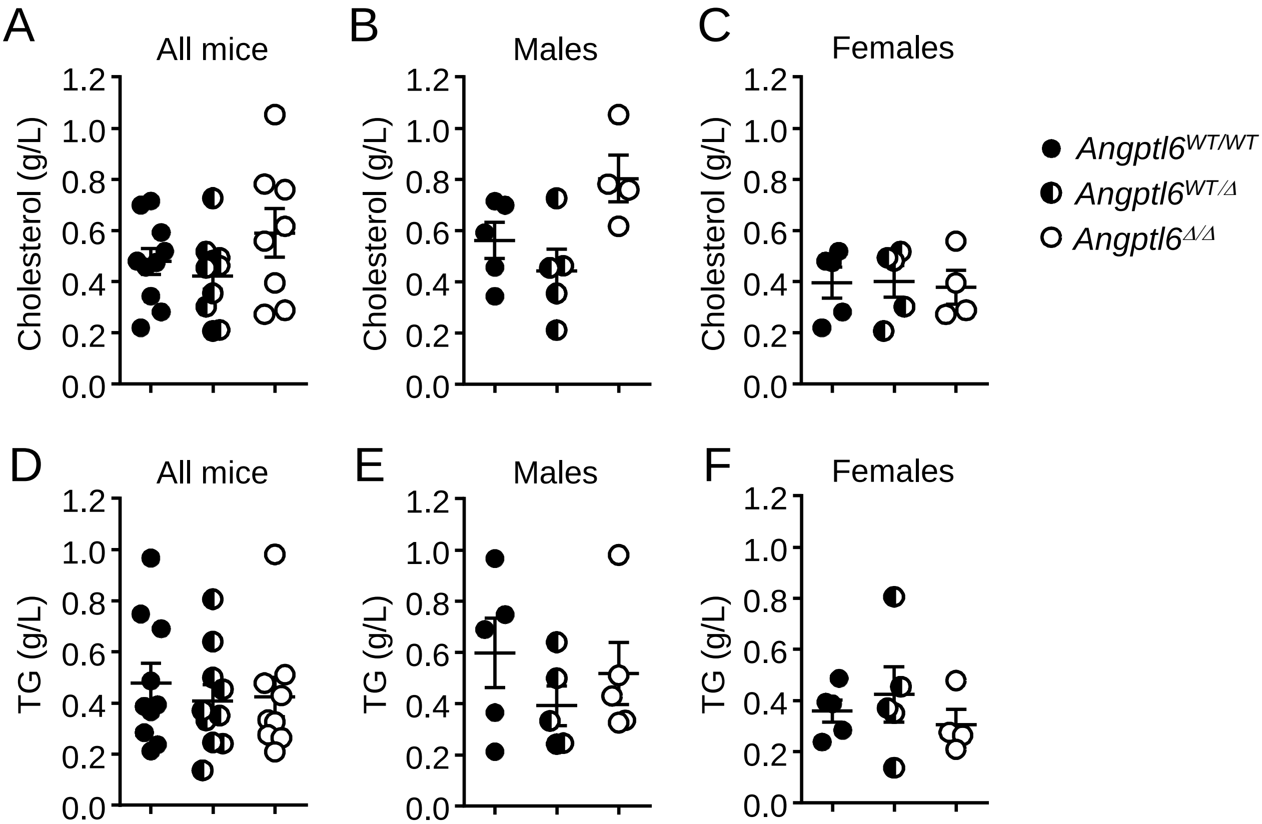


**Figure S2.** Plasma cholesterol and triglycerides (TG) concentration of *Angptl6*$^{WT/WT}$, *Angptl6*$^{WT/\Delta}$ and *Angptl6*$^{\Delta/\Delta}$ mice. **A-C**, Plasma cholesterol in all mice (A), male mice (B) and female mice (C). **D-F**, TG in all mice (D), male mice (E) and female mice (F). (*ns* between genotype and sex, One -way ANOVA, Bonferroni post-hoc test).

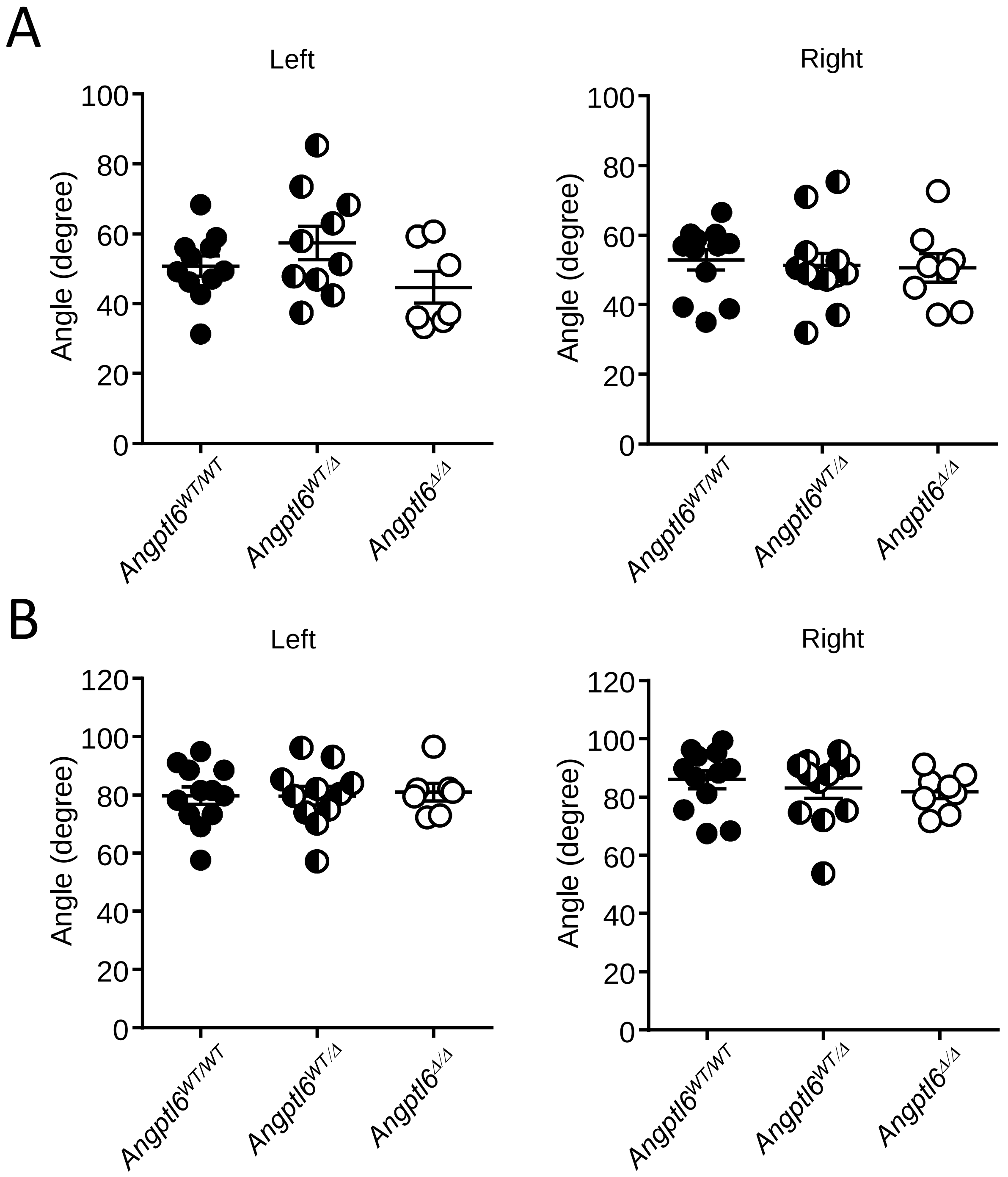


**Figure S3.** Angles of left and right arterial bifurcations of the circle of Willis of *Angptl6*$^{WT/WT}$, *Angptl6*$^{WT/\Delta}$ and *Angptl6*$^{\Delta/\Delta}$ mice. **A,** Anterior cerebral artery-olfactive artery bifurcation. **B,** Internal carotid artery-middle cerebral artery bifurcation. (*ns* between genotype, One -way ANOVA, Bonferroni post-hoc test).

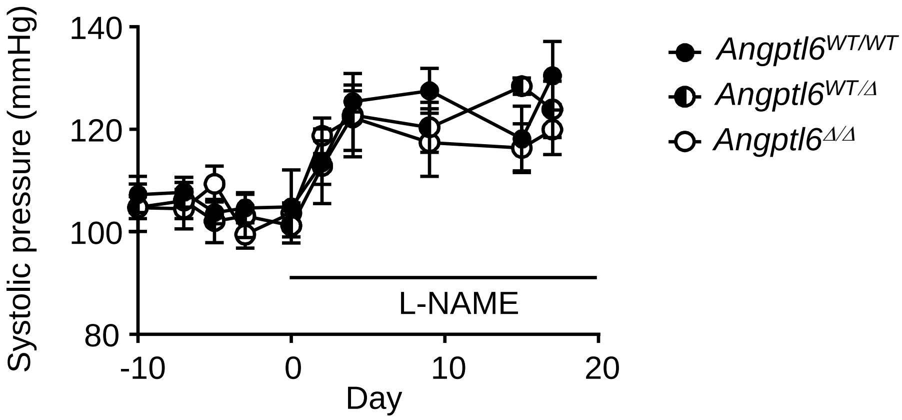


**Figure S4.** Systolic blood pressure in *Angptl6*$^{WT/WT}$, *Angptl6*$^{WT/\Delta}$ and *Angptl6*$^{\Delta/\Delta}$ mice before and during L-NAME administration. (Data shown are the mean ± s.e.m. of 3-7 mice; 2-way ANOVA, *ns* between genotype).

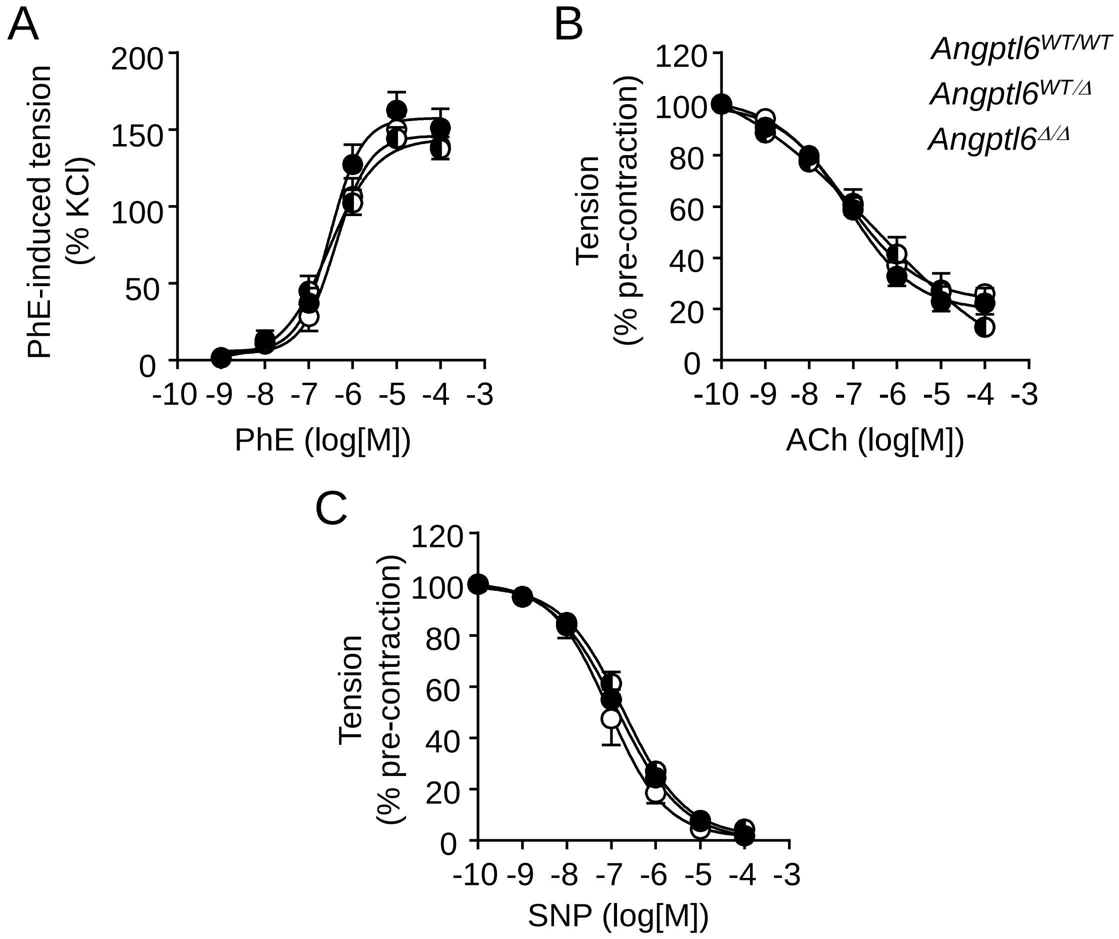


**Figure S5.** Contractile and relaxation properties of mesenteric arteries of *Angptl6*$^{WT/WT}$, *Angptl6*$^{WT/\Delta}$ and *Angptl6*$^{\Delta/\Delta}$ mice. A, Cumulative concentration-response curves for the contraction of mesenteric artery rings induced by PhE. Contraction is expressed as a percentage of the maximal KCl (90 mM)-induced contraction. B and C, Cumulative concentration-response curves for the relaxation of mesenteric artery rings induced by ACh (B) and SNP (C). Tenion is expressed as a percentage of the maximal pre-tension induced by PhE/5-HT. (Data shown are the mean ± s.e.m. of 13-24 mice in A, 14-22 mice in B and 5-14 mice in C; 2-way ANOVA, *ns* between genotype).

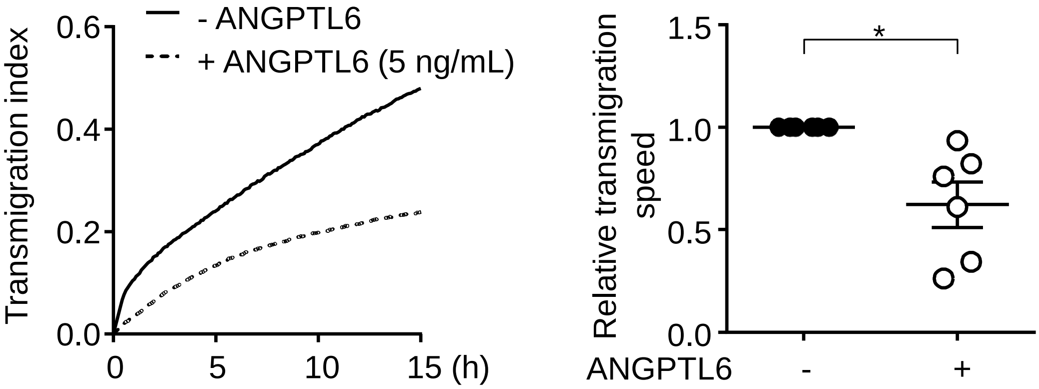


**Figure S6.** ANGPTL6 coating decreases human arterial smooth muscle cell migration. A. Typical curves showing the cell index representing the transmigration of human arterial smooth muscle cells across a membrane coated with gelatin alone (black line) and on ANGPTL6-containing gelatin (5 mg/mL, dotted line) over time. B. Quantification of the transmigration calculated from the slope of the experimental curves (*p<0.05; Wilcoxon test).
